# Single Cell Transcriptome Signatures of Sarcoidosis in Lung Immune Cell Populations

**DOI:** 10.1101/2025.01.20.633917

**Authors:** Camille Moore, Shu-Yi Liao, Cheyret Wood, Arunangshu Sarkar, Jonathan Cardwell, Kristyn MacPhail, Margaret M. Mroz, Christina Riley, Kara Mould, Clara Restrepo, Li Li, Lisa A. Maier, Ivana V. Yang

**Author notes:** contributed equally, co-first author (Camille Moore, Shu-Yi Liao). contributed equally, co-senior author (Lisa A. Maier, Ivana V. Yang).

## Abstract

**Rationale:** To identify cell specific molecular changes associated with sarcoidosis risk and progression, we aimed to characterize the cellular composition, gene expression patterns, and cell-cell interactions in BAL cells from patients with sarcoidosis (both progressive and non-progressive) and healthy controls.

**Methods:** Single cell RNA-seq data were collected on 12 sarcoidosis and 4 control participants. We combined scRNA-seq data from these participants with our previously collected data on 4 sarcoidosis and 10 control participants for a final sample size of 16 sarcoidosis cases (8 progressive and 8 non-progressive) and 14 controls. Following initial preprocessing in CellRanger, data were quality controlled, combined, and clustered in Seurat. We tested differences in cell proportions by disease group using F-tests on cell proportions and differences in gene expression using pseudobulk analysis. Cell to cell communication and pathway analysis were performed using CellChat.

**Results:** We identified five macrophage populations: resident, high metallothionein (MT) resident, recruited, profibrotic recruited, and proliferating macrophages. Each subpopulation displayed unique gene expression profiles, with notable differential expression of genes and pathways linked to sarcoidosis in resident macrophages, recruited macrophages, and proliferating macrophages. We also observed changes in gene expression associated with disease progression in resident and recruited macrophages. In non-macrophages cells, we observed a significant reduction in the number of B cells in sarcoidosis patients compared to controls. Among T cell populations, we identified specific transcriptional alterations at gene and pathway level. Additionally, we observed distinct differences in cell-to-cell interactions of macrophages and T cells between sarcoidosis patients and healthy controls.

**Conclusions:** These findings underscore the complexity of immune cell involvement in sarcoidosis and highlight potential cellular and molecular targets for further investigation.

## INTRODUCTION

Sarcoidosis is a complex, multi-system granulomatous disease characterized by immune dysregulation and chronic inflammation, primarily affecting the lungs(1). Its exact etiology remains unclear; however, it is believed to result from an exaggerated immune response to environmental or infectious triggers in genetically predisposed individuals. The hallmark of sarcoidosis is the formation of non-necrotizing granulomas, primarily composed of activated macrophages and T lymphocytes (2), contributing to tissue damage and in some fibrosis. Despite its heterogeneous presentation, distinguishing between progressive and non-progressive or resolving disease remains challenging, emphasizing the need for better diagnostic biomarkers and therapeutic targets (3).

Recent advances in single-cell RNA sequencing (scRNA-seq) have provided an unprecedented level of detail in understanding the cellular landscape and transcriptional profiles associated with sarcoidosis. For example, a recent study used scRNA-seq together with spatial transcriptomics to identify an immune stimulatory environment in granulomas that repurposes transcriptional programs associated with lymphoid organ development (4). This work highlighted the role of monocytes in driving granuloma formation in skin and lung. However, the only study that has addressed disease risk and progression in lung in bronchoalveolar lavage (BAL) cells was our pilot study with a limited number of samples (5). Leveraging this knowledge gap, we aimed to better characterize the cellular composition and gene expression patterns in BAL cells from patients with sarcoidosis (both progressive and non-progressive) and healthy controls. We integrated our current dataset with our previously published data (5, 6) to increase statistical power. By focusing on immune cell subsets, particularly macrophages and T cells, we sought to unravel the molecular mechanisms driving disease progression and identify novel therapeutic targets.

## METHODS

### Sample Selection and Single Cell RNA-Seq Data Collection

#### Sample Sources

Sarcoidosis cases and non-diseased controls were enrolled at National Jewish Health. 12 sarcoidosis and 4 control participants were enrolled for the purpose of the current study. We combined scRNA-seq data from these participants with our previously collected data on 4 sarcoidosis (GSE184735) and 10 control (GSE151928) participants for a final sample size of 16 sarcoidosis cases and 14 controls. All sarcoidosis cases were phenotyped using the same criteria (7). Bronchoscopies, BAL cell processing, and scRNA-seq data collection were performed using uniform protocols for samples from our current and previous studies.

#### Clinical Phenotyping

For all sarcoidosis cases, medical records were reviewed for clinical features including acuity of presentation (acute/non-acute), organ involvement, pulmonary function tests (PFT), chest imaging and immunosuppressive treatment at time of BAL and up to two years after BAL (7). All sarcoidosis subjects met the ATS/ERS criteria(8) for tissue biopsy confirmation of diagnosis of sarcoidosis. The non-progressive phenotype was defined as having either acute (i.e. consistent with Lofgren’s syndrome) or non-acute disease presentation, no new organ involvement, lung function testing with <10% decline in FVC or FEV_1_, <15% decline in DL_CO,_ and stable chest imaging within 2 years after BAL. The progressive phenotype had a non-acute disease presentation; lung function testing with ≥10% decline in FVC or FEV_1_; or ≥15% decline in DL_CO_; worsening chest imaging; and/or if they required initiation of systemic immunosuppressive treatment any time up to 2 years after BAL.

#### Bronchoalveolar lavage (BAL)

Bronchoscopy with BAL was performed as previously described.(9, 10). Briefly, four to eight 60 ml aliquots of sterile normal saline at room temperature were instilled into the airway with a syringe then aspirated using low suction. For each collection from an individual patient, the aspirated BAL specimens were pooled, kept on ice, and processed within one hour of collection. The BAL cells were obtained by centrifugation and cryopreserved.

#### Single-cell RNA-Seq Data Collection

Cryopreserved cells were thawed and captured on 10X Chromium in the Genomics Core at National Jewish Health using 10x Genomics Single Cell 3′ v3 chemistry, followed by sequencing on Illumina NovaSeq.

### Single-cell RNA-Seq Computational Pipeline and Preliminary Quality Control

Initial pre-processing of the 10x Genomics scRNA-Seq data from 30 participants, including demultiplexing, alignment to the hg38 human genome, and unique molecular identifier–based (UMI-based) gene expression quantification, was performed using Cell Ranger (Mould samples: version 3.0.2, this study and Liao samples: version 3.1.0, 10x Genomics). Data were available on 171,003 cells from 30 samples. We filtered out low-quality cells with fewer than 100 genes detected or with greater than 25% of mapped reads originating from the mitochondrial genome. We additionally safeguarded against doublets by removing cells with a UMI count greater than the 98th percentile of UMI counts for each sample. Prior to downstream analysis, select mitochondrial and ribosomal genes (genes beginning with MT-, MRPL, MRPS, RPL, or RPS) were removed. The preliminary quality-controlled data set consisted of 150,176 cells and 20,491 genes. To account for differences in coverage across cells, we normalized and variance stabilized UMI counts using the SCTransform method in the Seurat version 5.1.0 R package. In addition to adjusting for sequencing depth, we also adjusted for the proportion of mitochondrial reads.

### Data Integration, Dimensionality Reduction, and Clustering

Data from the 30 participants were combined using single-cell integration implemented in Seurat v3 utilizing reference samples (4 from this study, 2 from Mould, and 1 from Liao). Integration was carried out using the top 30 dimensions from a canonical correlation analysis based on SCTransform-normalized expression of the top 3,000 most informative genes, defined by gene dispersion using the Seurat’s SelectIntegrationFeatures function. Integrated data were then clustered and visualized using the top 26 principal components. For visualization, we reduced variation to 2 dimensions using UMAP (n.neighbors = 50, min.dist = 0.3). Unsupervised clustering was performed by using a shared nearest neighbor graph based on 50 nearest neighbors and then by determining the number and composition of clusters using a smart local moving algorithm (resolution = 0.6). This algorithm identified 26 preliminary clusters.

### Cluster Markers

To identify cluster markers, we carried out pairwise differential expression analysis comparing log-normalized expression in each cluster to all others using a Wilcoxon rank sum test. Markers were identified as genes exhibiting significant upregulation when compared against all other clusters, defined by having a Bonferroni-adjusted P < 0.05, a log fold change > 0.25, and >10% of cells with detectable expression. This analysis was then performed separately for each participant using Seurat’s FindConservedMarkers function to determine if marker genes were consistent across participants. Using these markers, 23 clusters were identified as either macrophage or nonmacrophage cells. The remaining three clusters were unable to be identified and were excluded; all three had low number of cells (≤330), one had high number of mitochondrial reads, the other two had high amounts of ribosomal RNA. The dataset was divided into macrophage (72,935 cells) and nonmacrophage (23,806 cells) data sets for further analysis.

### Additional Quality Control and Re-clustering

Dimensionality reduction and clustering were performed separately for the macrophage and non-macrophage datasets as described above, resulting in 17 macrophage and 16 nonmacrophage clusters. Potential doublets were assessed with the scds R package, which calculates a hybrid of the co-expression based and binary classification doublet scores to for each cell. Cells with high hybrid doublet scores (hybrid > 0.5) were excluded from further analysis in the macrophage dataset (this step was not necessary in non-macrophage cells). We also examined marker expression and excluded macrophage cluster 11 based on low expression of *CD68, FABP4, FCN1, and VCAN*. Similarly, non-macrophage clusters 10 and 14 were determined to be composed of a mixture cells, so these cells were excluded from further analysis. This resulted in 71,616 cells in the macrophage and 22,810 cells in the non-macrophage datasets.

### Cell Proportion Analysis

We tested differences in cell proportions by disease group using F-tests on cell proportions calculated by the propeller R package. Propeller calculates cell type proportions for each biological replicate, performs a variance stabilizing transformation (logit) on the matrix of proportions, and fits a linear model for each cell type or cluster using the limma framework.

### Pseudo-bulk Differential Expression

To identify differentially expressed genes (DEGs) between control and sarcoidosis samples while accounting for clustering of cells within participants, we performed pseudo-bulk differential expression analysis separately for each cell type. Within each cell type, expression counts were summed across all cells for a participant, resulting in a single expression measurement for each gene for each participant. Low expressing genes were removed as defined of having less than 1 count per million in 1/3 of the samples. Pseudo-bulk expression was compared between participants with sarcoidosis and controls using bulk RNA-Seq analysis methods with the DESeq2 R package. Genes with Benjamini-Hochberg adjusted P values less than 0.05 were considered differentially expressed. This analysis was repeated to identify genes differentially expressed between progressive sarcoidosis and non-progressive sarcoidosis.

### Cell-level Differential Expression

Linear mixed models were used to compare cell level gene expression between sarcoidosis and control for each cell type. Genes were included in the analysis if they were expressed in at least 1% of the cells. DESeq2’s variance stabilizing transformation (VST) was used to normalize the UMI count data prior to modeling. The linear mixed model was fit with diagnosis (sarcoidosis or control) as a fixed effect and a random intercept for each subject to account for the clustering of cells within each subject.

### Cell-cell Interaction Analysis

Cell to cell communication and pathway analyses were performed using CellChat version 2.1.2. Communication was estimated separately for the sarcoidosis samples and controls samples for the following cell types; CD4+ T-cells, CD8+ T-cells, Resident Macrophages, High MT Resident Macrophages, Recruited Macrophages, Pro-fibrotic Recruited Macrophages, and Proliferating Macrophages. The human CellChat database was used to define the ligand-receptor interaction pairs and to group these pairs into functionally related pathways. Average gene expression was computed using the 25% truncated mean. 10,000 permutations were used to estimate cell communication probability. We filtered out communications that were not present in at least 75% of the samples. To account for multiple testing of the pathways, statistically significant communication was defined as having a Bonferroni corrected p-values less than 0.05. Interaction strength and number of interactions between cell types were compared between the sarcoidosis and control groups.

## RESULTS

### Enrolled Participants

Our single cell RNA-Seq dataset consisted of BAL cells from 30 participants: 16 sarcoidosis (8 progressive and 8 non-progressive) and 14 non-diseased controls. Characteristics of the participants whose samples were included in the study are summarized in **Table 1**. Sarcoidosis patients were older than controls and included some former smokers, while there were no former smokers in the control group. 14 of these samples were from our previous studies; this includes 4 sarcoidosis (2 progressive and 2 non-progressive) (5) and 10 control (6) samples collected using the same protocols for sarcoidosis phenotyping, BAL collection, and scRNA-seq data collection at National Jewish Health.

**Table 1.**
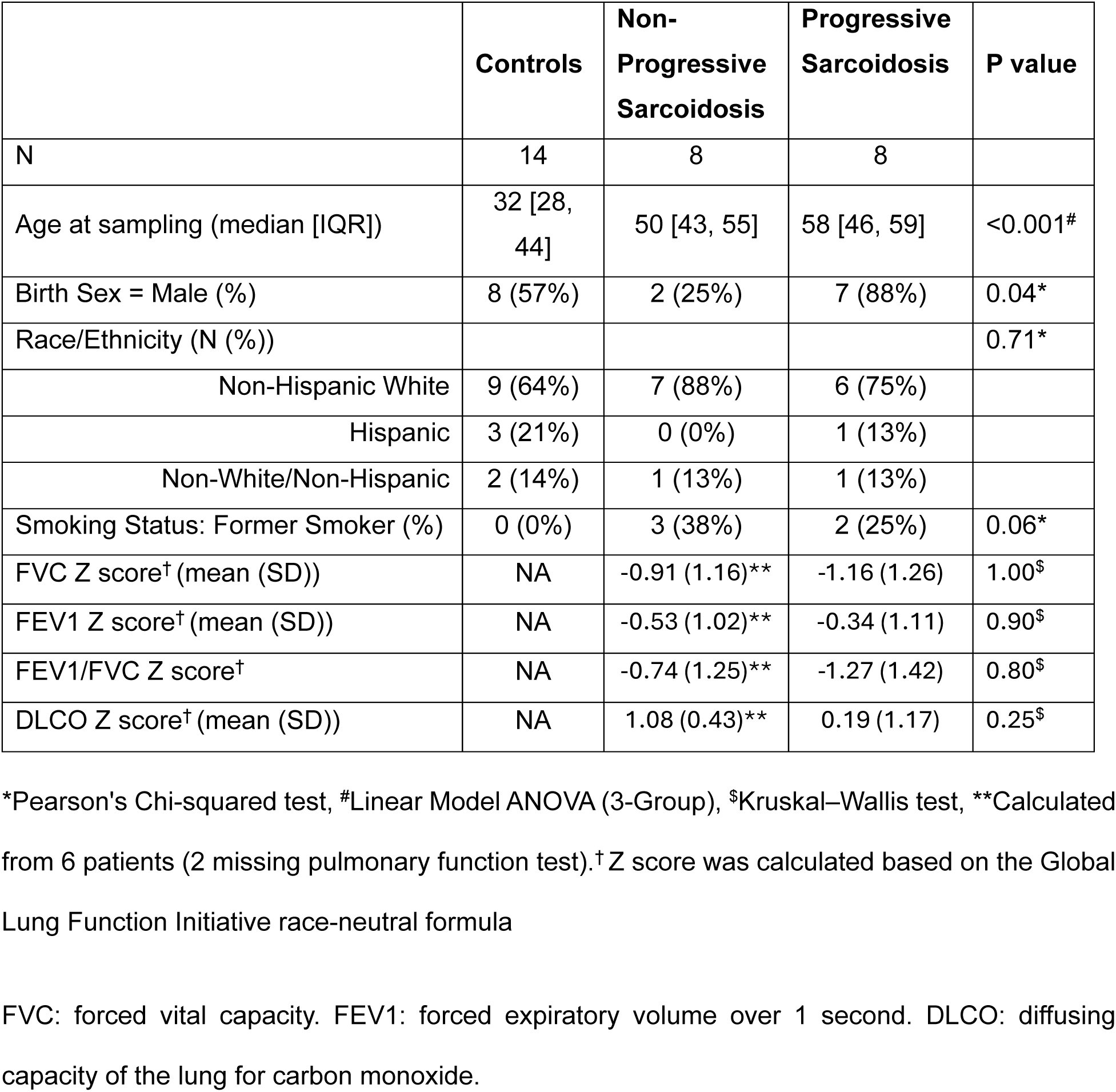
Demographic and clinical characteristics of the control, progressive sarcoidosis, and non-progressive sarcoidosis subjects.

|  | **Controls** | **Non-Progressive Sarcoidosis** | **Progressive Sarcoidosis** | **P value** |
| --- | --- | --- | --- | --- |
| N | 14 | 8 | 8 |  |
| Age at sampling (median [IQR]) | 32 [28, 44] | 50 [43, 55] | 58 [46, 59] | <0.001<sup>#</sup> |
| Birth Sex = Male (%) | 8 (57%) | 2 (25%) | 7 (88%) | 0.04* |
| Race/Ethnicity (N (%)) |  |  |  | 0.71* |
| Non-Hispanic White | 9 (64%) | 7 (88%) | 6 (75%) |  |
| Hispanic | 3 (21%) | 0 (0%) | 1 (13%) |  |
| Non-White/Non-Hispanic | 2 (14%) | 1 (13%) | 1 (13%) |  |
| Smoking Status: Former Smoker (%) | 0 (0%) | 3 (38%) | 2 (25%) | 0.06* |
| FVC Z score<sup>†</sup> (mean (SD)) | NA | -0.91 (1.16)** | -1.16 (1.26) | 1.00<sup>\$</sup> |
| FEV1 Z score<sup>†</sup> (mean (SD)) | NA | -0.53 (1.02)** | -0.34 (1.11) | 0.90<sup>\$</sup> |
| FEV1/FVC Z score<sup>†</sup> | NA | -0.74 (1.25)** | -1.27 (1.42) | 0.80<sup>\$</sup> |
| DLCO Z score<sup>†</sup> (mean (SD)) | NA | 1.08 (0.43)** | 0.19 (1.17) | 0.25<sup>\$</sup> |

\*Pearson's Chi-squared test, <sup>#</sup>Linear Model ANOVA (3-Group), <sup>\$</sup>Kruskal–Wallis test, \*\*Calculated from 6 patients (2 missing pulmonary function test).<sup>†</sup> Z score was calculated based on the Global Lung Function Initiative race-neutral formula
FVC: forced vital capacity. FEV1: forced expiratory volume over 1 second. DLCO: diffusing capacity of the lung for carbon monoxide.

### Transcriptional Signatures of Disease Risk in Macrophage Cell Populations

Following quality control and clustering described in Methods, we split our dataset into macrophage and non-macrophage cells to enable more refined clustering. The macrophage dataset comprised resident macrophages (64,143 cells; high expression of *FABP4, CD68, MRC1*), of which 2,764 were marked by high metallothionein (MT) gene expression; recruited monocyte-derived macrophages (6,474 cells; high expression of *FCN1, VCAN, CCL2*), of which 1,263 were pro-fibrotic recruited macrophages (high expression of *CHI3L1, SPP1, MMP9, MARCKS, PLA2G7*); and proliferating macrophages (999 cells; high expression of *MKI67, MCM2, PCNA*) (**Figure 1A**). We first examined the proportion of these five macrophage populations across the samples (**Figure 1B**) and determined that there were no significant differences in sarcoidosis samples compared to controls (**Supplemental Table S1A**). Differential gene expression analysis on pseudobulk counts identified significant differences (FDR-adjusted p-value<0.05) associated with sarcoidosis in 16 genes in resident macrophages, 3 in high MT resident macrophages, 28 in recruited, 2 in profibrotic recruited, and 1 in proliferating macrophage populations (**Supplemental Table S2**). We also tested for differences in cell-level gene expression using linear mixed-effects models. While that analysis did not result in significant differences at the stringent FDR-adjusted p<0.05, top genes were similar to those identified by pseudobulk testing (**Supplemental Table S3**). Representative violin plots for significantly DEGs by pseudobulk analysis are shown in **Figure 1C**. Among transcripts associated with sarcoidosis, we identified increased gene expression in resident macrophages in interleukin 1 receptor type (*IL1R1*), receptor for IL-1α, IL-1bβ, and interleukin-1 receptor antagonist (ILRAP); a novel macrophage actin protein proline-serine-threonine phosphatase interacting protein 2 (*PSTPIP2*), and TAP binding protein (*TAPBP*) that is essential for optimal peptide loading on the MHC class I molecule.

**Figure 1.**
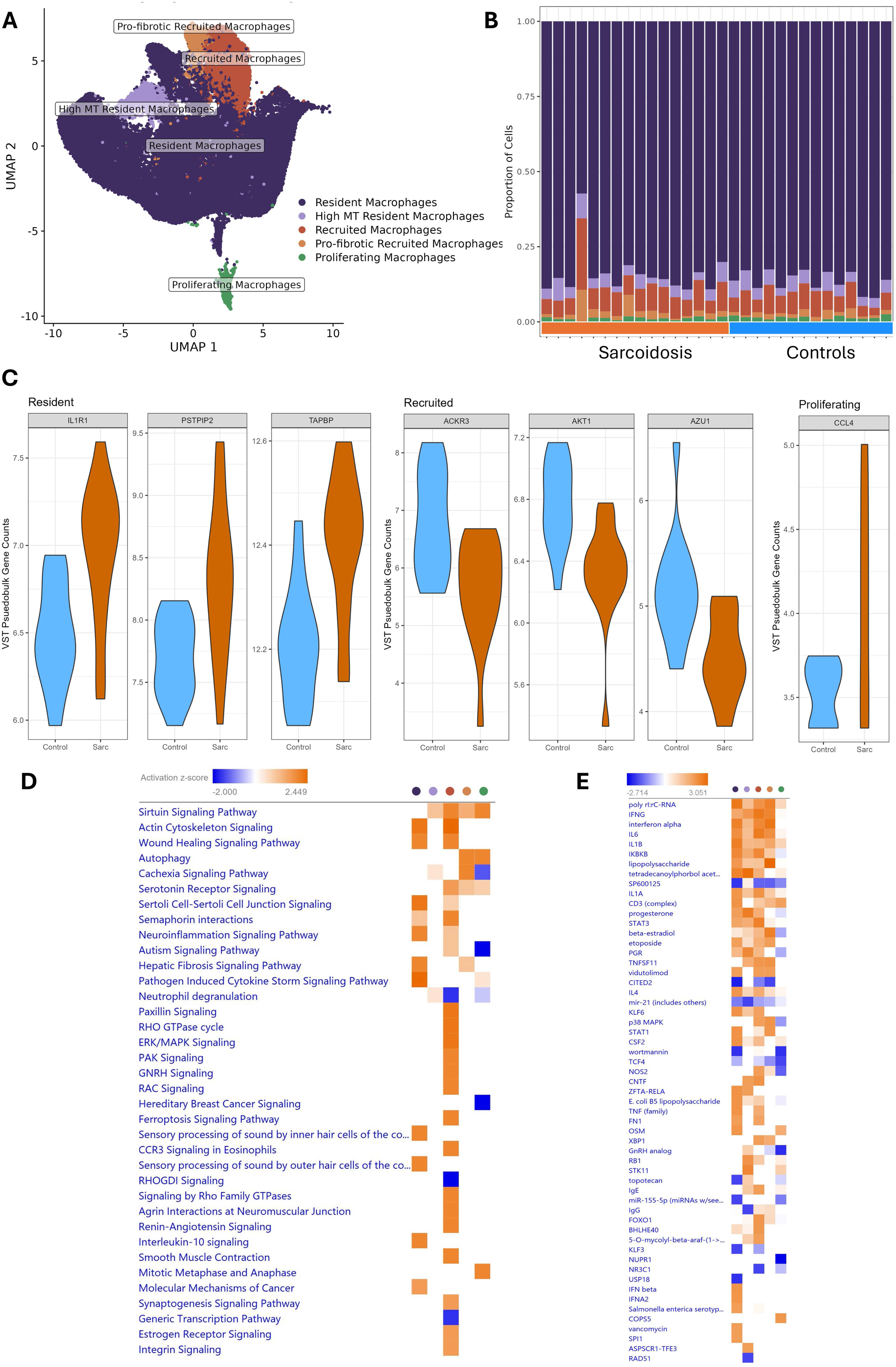
Sarcoidosis transcriptome signatures in macrophage cell populations. **(A)** UMAP projections of five macrophage clusters. **(B)** Proportion of cells in each of the five macrophage clusters across 30 individuals included in the analysis. **(C)** Violin plots of select genes differentially expressed in sarcoidosis (N = 16) compared to controls (N = 14) in specific macrophage populations. **(D/E)** Ingenuity pathway (D) and upstream regulator (E) analysis of top 100 DEGs in sarcoidosis compared to controls. Colors in Panels D and E indicate cell populations using the same color legend as the UMAP in Panel A.

In recruited macrophages, three downregulated transcripts, atypical chemokine receptor 3 (*ACKR3*), AKT serine/threonine kinase 1 (*AKT1*), and azurocidin 1 (*AZU1*) were among the DEGs. Furthermore, we highlight upregulation of C-C motif chemokine ligand 4 (*CCL4,* also known as *MIP1β*), ligand for the CCR5 receptor, in proliferating macrophages. To identify gene expression changes associated with sarcoidosis beyond the few highly statistically significant genes, we performed pathway and upstream regulator analysis of top 100 most significant genes (by nominal p value) in the five macrophage clusters. Generally, we observed cell population specific pathway activation with recruited macrophages having the most pronounced changes (**Figure 1D**). Among pathways activated in recruited macrophages were ERK/MAPK, RHO GTPase, CCR3, ferroptosis, and integrin signaling. On the other hand, resident macrophages were marked by activation of IL10 signaling. A few pathways were also activated across cell populations; including sirtuin signaling in all but resident macrophages, wound healing in resident and recruited macrophages, and autophagy in profibrotic and proliferating macrophages. Upstream regulator analysis identified activation of many shared and unique transcriptional regulators (**Figure 1E**). All but proliferating macrophages were marked by activation of cytokines IFNG, IL6, IL1, and IL4.

### Transcriptional Signatures of Disease Progression in Macrophage Cell Populations

Focusing on disease progression, we did not observe any significant changes in cell proportions across macrophage populations (**Supplemental Table S1B)**. Pseudobulk analysis identified 16 transcripts in resident macrophages significantly (FDR-adjusted p<0.05) differentially expressed in progressive compared to non-progressive sarcoidosis (**Figure 2A**); another 14 were significant at FDR-adjusted p<0.1 (**Supplemental Table S4**). Surprisingly, the majority of transcript levels for the majority of the significant genes were similar in progressive sarcoidosis and controls, with non-progressive cases demonstrating different levels compared to controls and progressive cases. Minimal significant changes (1-2 genes each) were identified in the remaining three macrophage populations, likely due to lower power to detect changes in these smaller cell clusters. Among transcripts upregulated in resident macrophages from participants with progressive disease was the MHC class II molecule *HLA-DRB5* (also upregulated in other macrophage populations), while downregulated transcripts include C-C motif chemokine receptor 5 (*CCR5)* and Interleukin 3 Receptor Subunit Alpha *(IL3RA)*. We compared pathways enriched in top 100 genes associated with sarcoidosis and progression in resident macrophages and found very few pathways dysregulated in progressive compared to non-progressive sarcoidosis; this is in contrast to pronounced dysregulation, mainly activation, we observed in sarcoidosis compared to controls (**Figure 2B**). We observed activation of Pathogen-induced cytokine storm and Costimulation by CD28 family pathways in both disease and progression analyses while pathways such as Phagosome formation and Neutrophil deregulation demonstrated opposite effects in the two comparisons (activation in disease but deactivation in progressive compared to non-progressive disease). Similarly, most upstream regulators were activated in disease but deactivated in progressive compared to non-progressive disease comparison (**Figure 2C**).

**Figure 2.**
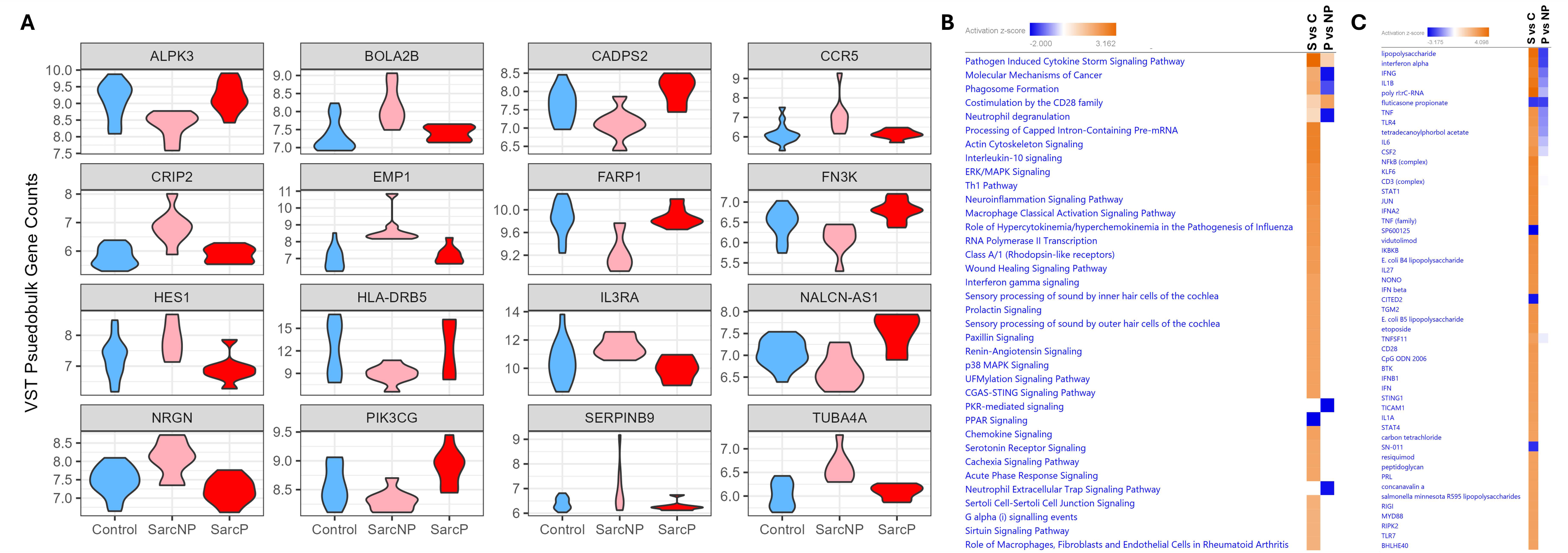
Sarcoidosis progression transcriptome signatures in macrophage cell populations. **(A)** Violin plots of 16 genes differentially expressed in resident macrophages of sarcoidosis compared to controls progressive compared to non-progressive sarcoidosis. **(B/C)** Ingenuity pathway (B) and upstream regulator (C) analysis of top 100 differentially expressed genes in progressive sarcoidosis (SarcP; N = 8) compared to non-progressive sarcoidosis (SarcNP; N = 8). S vs C refers to sarcoidosis compared to control analysis and P vs NP refers to SarcP compared to SarcNP analysis.

### Transcriptional Signatures of Disease Risk in Non-Macrophage Cell Populations

As expected, the majority of non-macrophage cells were comprised of T-cells (18,047 cells), of which 6,399 cells were CD4+ T-cells, 8,800 were CD8+ T-cells, and the remaining 2,848 were mixed T-cells. Other non-macrophage clusters included dendritic cells (DC; 2,597 cells), B-cells (823 cells), natural killer (NK) cells (757 cells), plasmacytoid dendritic cells (pDC; 154 cells), and epithelial cells (432) (**Figure 3A**). We first examined proportions of these cell populations across the samples (**Figure 3B**) and observed a significant decrease (p=0.01) in B cells in sarcoidosis (2.6% of non-macrophage cells) compared to controls (7.5% of non-macrophage cells) (**Supplemental Table S5**). Differential gene expression analysis on pseudobulk counts identified significant differences (FDR-adjusted p-value<0.05) associated with sarcoidosis in five transcripts in all T cells, 8 when subset to CD4+ T cells, and one in dendritic cells (**Supplemental Table S6**). Representative violin plots for significantly differentially expressed genes by pseudobulk analysis are shown in **Figure 3C**. Among transcripts associated with sarcoidosis, we highlight upregulation of unknown transcript *AC009053.2*, chemokine ligand 2 (*CCL2*) and transcription factor *TWIST1* in combined T cells, upregulation of acidic nuclear phosphoprotein 32 family member E (*ANP32E*) and clusterin (*CLU*) in CD4+ T cells, and downregulation all *AZU1* in all T cells and when focused to CD4+ T cells. Pathway analysis of top 100 transcripts (**Figure 3D**) highlighted a few shared and mostly unique pathways across the cell populations we examined; activation of T cell exhaustion pathway in CD8+ T cells is noteworthy. Most changes in upstream regulators were observed in CD4+ T cells, including but not limited to activation of TNF, IFNG, and IL1B; other cell populations demonstrated less prominent transcriptional changes (**Figure 3E**). Given the small sample sizes, we did not expect to identify many changes associated with disease progression. Analysis of disease progression in non-macrophage cell clusters identified only a few significant differences at FDR-adj<0.05 likely because of small number of cells and samples.

**Figure 3.**
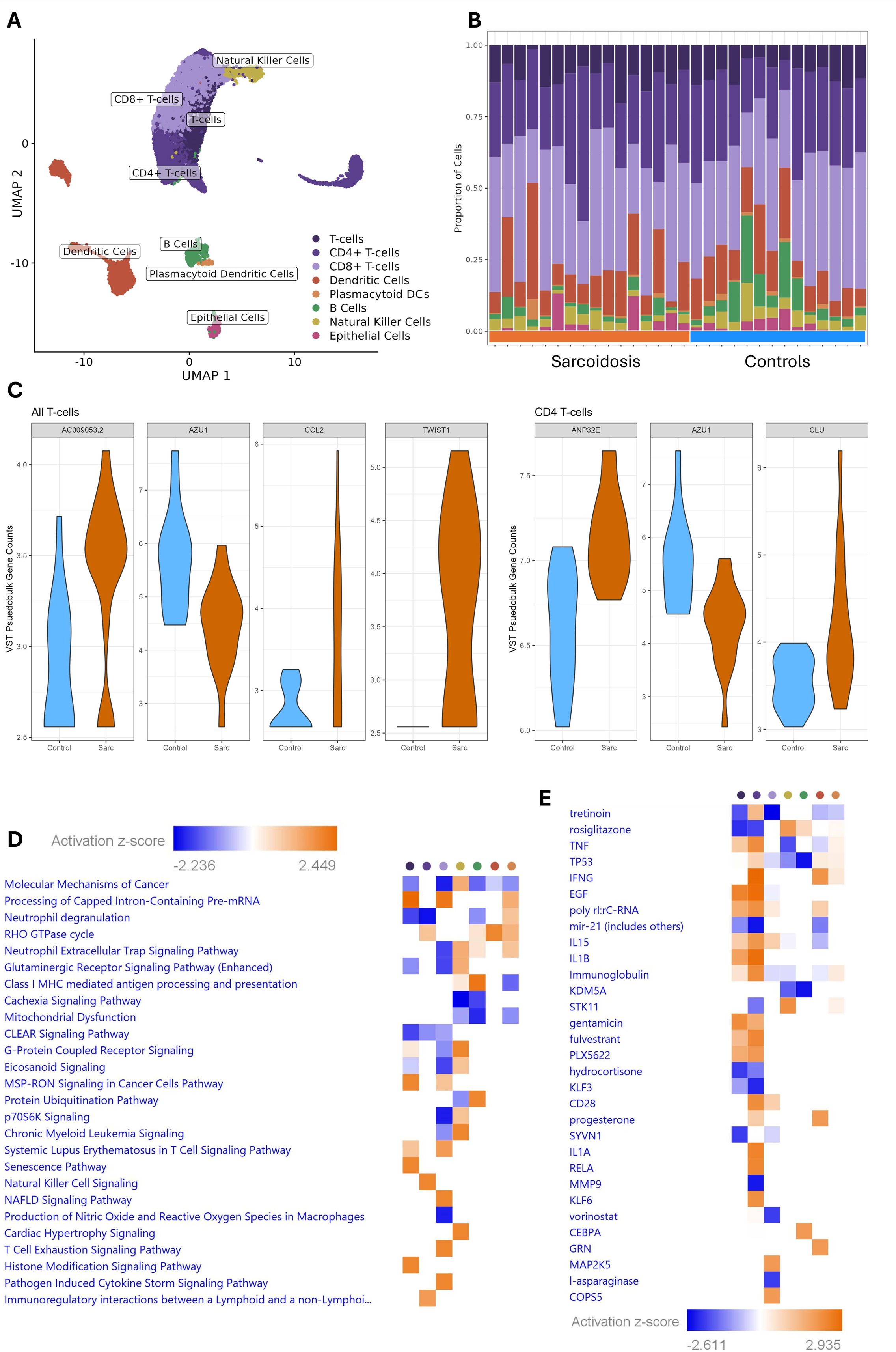
Sarcoidosis transcriptome signatures in non-macrophage cell populations. **(A)** UMAP projections of cell clusters. **(B)** Proportion of cells in each of the clusters across 30 individuals included in the analysis. **(C)** Violin plots of select genes differentially expressed in sarcoidosis (N = 16) compared to controls (N = 14) in specific cell populations. **(D/E)** Ingenuity pathway (D) and upstream regulator (E) analysis of top 100 differentially expressed genes in sarcoidosis compared to controls. Colors in Panels D and E indicate cell populations using the same color legend as the UMAP in Panel A.

### Cell-Cell Interaction Analysis of Macrophage and T Cell Populations

In the final set of investigations, we performed ligand-receptor analysis on sarcoidosis and control single cell datasets to study the crosstalk between macrophage and T cell populations (we excluded other cell types due to small numbers). Globally, we observed reduced signaling in sarcoidosis compared to controls (**Figure 4A**), with only one pathway (GAS signaling) exhibiting stronger communication in sarcoidosis and 25 pathways with stronger communication in controls; these include immune pathways such as CCL, CXCL, and MHC class II, as well as pro-fibrotic pathways such as ICAM, PECAM, and TGFβ. We observed similar patterns of reduced ligand-receptor signaling in sarcoidosis compared to controls at the cell type level (**Figure 4B**). We also observed a larger number of changes in receiving or sending signals in macrophage than T cell populations (**Supplemental Figure S1A**). Considering interactions across macrophage and T cell populations (**Figure 4C**), we observed a higher number of interactions and interaction strength among most cell populations in controls (blue lines), however, the number of interactions and interaction strength of CD4+ T cells with other cell types were higher in sarcoidosis (red lines). We then focused on cell-specific ligand-receptor analysis to identify dysfunctional signaling in sarcoidosis compared to controls. Comparing the communication probabilities, we observed many interactions decreased and downregulated in sarcoidosis, and a few increased and upregulated in sarcoidosis (**Supplemental Figure S2**). Focusing on only interactions between macrophage and T cell populations, we observed differences in interactions of all macrophage populations with CD4+ and CD8+ T cells (**Figure 4D**, **Supplemental Figure S1B**). This includes upregulation of FN1-CD44 signaling from high MT resident and proliferating macrophages to CD4+ and CD8+ T cells as well as downregulation of LGALS9-CD45 signaling from all macrophage populations to CD4+ and CD8+ T cells. When looking at T cell to macrophage signaling, more specific downregulation of HLA-DRB1-CD4 and HLA-F-LILRB1 signaling was observed from CD4+ and CD8+ T cells to high MT resident and profibrotic recruited macrophages. On the other hand, we observed upregulation of HLA-DRA-CD4, HLA-DPA1-CD4, and HLA-DPB1-CD4 signaling from CD4+ and CD8+ T cells to high MT resident and profibrotic recruited macrophages. These results, while initially surprising in directionality, are consistent with expression patterns of these molecules in macrophage and T cell populations we examined (**Supplemental Figure S3**).

**Figure 4.**
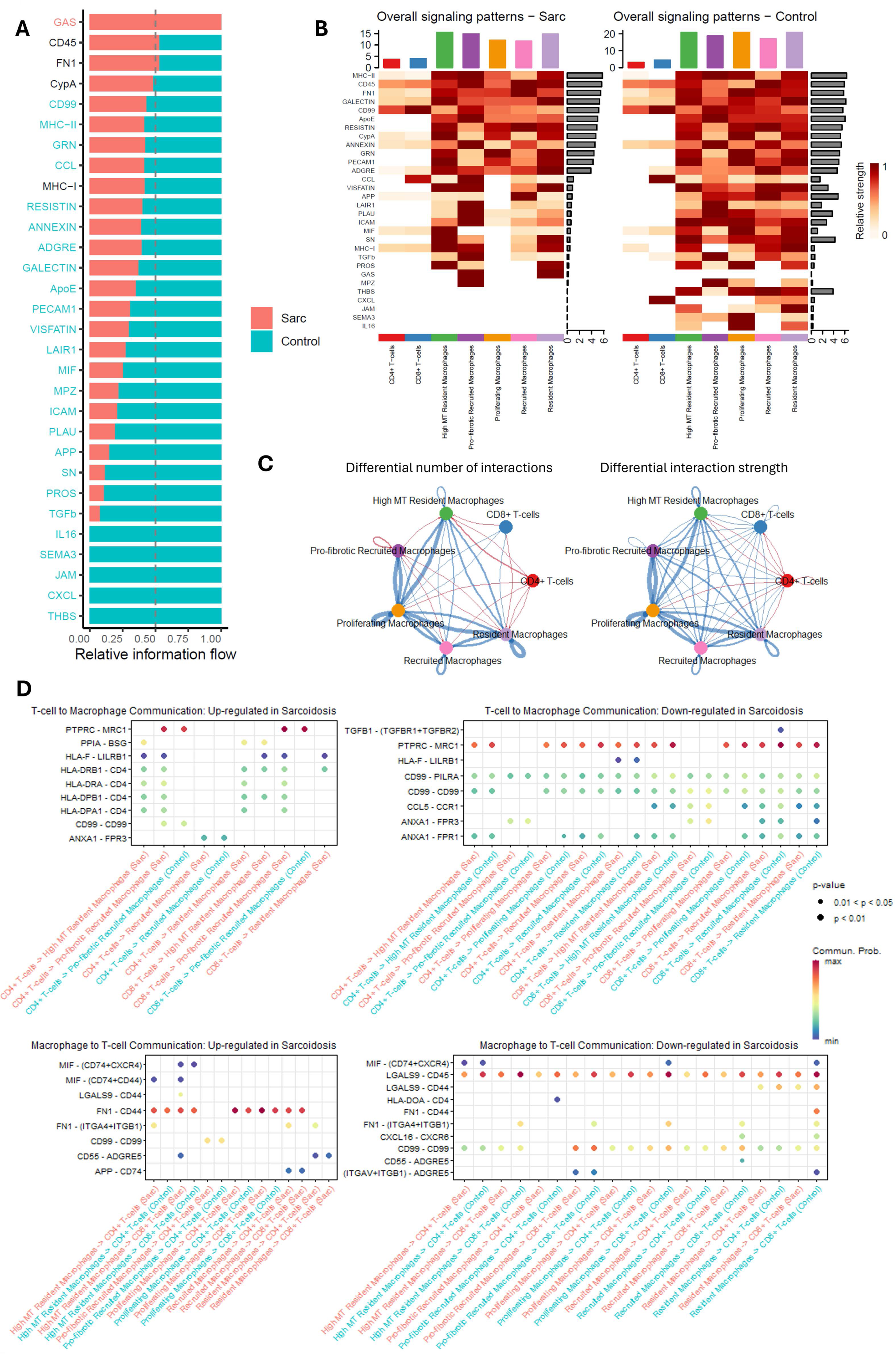
Cell-cell interaction analysis of macrophage and T cell populations. **(A)** Global analysis of ligand-receptor pathways between sarcoidosis and controls depicted as the colored bar plot comparing information flow for each pathway. Pathway labels are colored according to significant difference in signaling information, with enrichment in sarcoidosis (salmon) or controls (light blue). Black labels indicate no significant difference in signaling information between sarcoidosis and control. **(B)** Heatmaps for signaling in ligand-receptor pathway at the cell type level sarcoidosis (left) and controls (right). **(C)** Circle plots of differential number of interactions of interaction strength among cell populations. Blue lines indicate enrichment in interactions in controls and red in sarcoidosis. Thickness of the line indicates the strength of the differential enrichment between sarcoidosis and controls. **(D)** Comparison of specific ligand-receptor interactions in macrophage-T cells populations that are down- (left) and up-regulated (right) in sarcoidosis.

## DISCUSSION

Our single-cell RNA-Seq analysis of BAL cells from sarcoidosis patients and healthy controls revealed distinct macrophage subpopulations and gene expression patterns associated with sarcoidosis and disease progression. Specifically, we identified five macrophage populations: resident, high metallothionein (MT) resident, recruited, profibrotic recruited, and proliferating macrophages. Each subpopulation displayed unique gene expression profiles, with notable differential expression of genes and pathways linked to sarcoidosis in resident macrophages, recruited macrophages, and proliferating macrophages. We also observed changes in gene expression associated with disease progression in resident and recruited macrophages. In non-macrophages cells, we observed a significant reduction in the number of B cells in sarcoidosis patients compared to controls. Among T cell populations, we identified specific transcriptional alterations at gene and pathway level. Additionally, we observed distinct differences in cell-to-cell interactions of macrophages and T cells between sarcoidosis patients and healthy controls. These findings underscore the complexity of immune cell involvement in sarcoidosis and highlight potential cellular and molecular targets for further investigation.

Resident macrophages expressed several upregulated genes in sarcoidosis, such as *IL1R1*, *PSTPIP2*, and *TAPBP*, implicated in sarcoidosis pathology. In pulmonary sarcoidosis, IL-1 signaling helps recruit additional immune cells to granulomas, contributing to chronic inflammation that defines disease persistence (11). IL-1 was also identified in the upstream regulator analysis in all but proliferating macrophage populations. IL1R acts as a receptor for both IL-1β and IL-1 receptor antagonist, (IRAP), which are both upregulated in sarcoidosis BAL cells (IL-1β secretion in BAL is also increased) (12, 13). Importantly, IL-1β expression is increased in sarcoid granulomas and has been shown to be critical for granuloma formation in the trehalose 6,6’-dimycolate-granuloma mouse model (12) and IRAP has also been localized to the sarcoid granuloma in lung tissue (13). *PSTPIP2* and *TAPBP* have not been reported in previous sarcoidosis literature but are worth considering as important novel targets. PSTPIP2 is a negative regulator that plays key roles in macrophage activation, neutrophil migration, cytokine production, and osteoclast differentiation, and is linked to innate immune and autoinflammatory diseases (14). TAPBP is involved in antigen processing and presentation pathway and while genetic variation in TAP genes have not been associated with sarcoidosis (15, 16), antigen processing and presentation pathways have been associated with sarcoidosis (17) and beryllium-induced lung disease (18–20).

In sarcoidosis, recruited macrophages display downregulated genes such as *AKT1*, *ACKR3*, and *AZU1*. PI3K/AKT1 pathway plays a key role in sarcoidosis, influencing T cell function, with its deregulation linked to disease progression (7, 17). The PI3K/AKT1 pathway has also been implicated in activation of mTOR (17), which has been associated with granuloma formation (21). Previous studies demonstrated both reduced proliferative response and exhaustion of T cells in progressive sarcoidosis is thought to be driven in part by inhibition of PI3K/AKT1 signaling (22). While AKT1 has been previously associated with sarcoidosis, ACKR3 and AZU1 represent novel findings in the regulation of inflammation. ACKR3 (also known as CXCR7) plays a key role in inflammation and granuloma formation by regulating chemokine availability, including CXCL12 and CXCL11, which are crucial for immune cell recruitment (23, 24); CXCL11 is upregulated in serum of patients with sarcoidosis (25) and CXCL12/CXCR4 is increased in sarcoidosis granulomas (26). ACKR3 enhances inflammation by promoting pro-inflammatory leukocyte phenotypes and angiogenesis, while acting as a chemokine scavenger to maintain immune balance (27). ACKR3’s ability to modulate chemokine gradients suggests its influence on granuloma architecture (28). AZU1, an antimicrobial protein could indicate an impaired immune response to microbial antigens. While AZU1 is recognized for its involvement in antimicrobial activities and inflammatory processes, its specific role in sarcoidosis, particularly concerning impaired immune responses to microbial antigens proposed as triggers, has not been well-defined in the literature. The novelty of these dysregulated genes in the context of sarcoidosis underscores a complex regulation of immune activity, suggesting these pathways may provide new targets for therapeutic intervention.

Proliferating macrophages in sarcoidosis show an upregulation of *CCL4*. CCL4 is a chemokine that binds to CCR5 and recruits additional immune cells to the site of inflammation. CCL4 was previously shown as upregulated in the BAL of sarcoidosis patients(29) and a mouse model of pulmonary granulomatosis(30). Moreover, CCL4 has been shown to function as a chemokine neoantigen that induces beryllium-specific CD4+ T cells in CBD (31). Its receptor CCR5 is also upregulated in sarcoidosis (32) and genetic variants in *CCR5* (*33*) have been associated with sarcoidosis susceptibility.

Progressive and non-progressive sarcoidosis exhibit distinct gene expression patterns in resident and recruited macrophage populations, which can provide valuable insights into the disease’s trajectory and severity. Some of the genes we identified have an established role in sarcoidosis. In addition to being associated with disease susceptibility, as discussed above, *CCR5* genetic variation has also been associated with persistent lung involvement in sarcoidosis (34), We now identify expression of this gene as associated with disease progression specifically in resident macrophages. Similarly, *HLA-DRB5,* whose expression is increased in progressive compared to non-progressive sarcoidosis in all macrophage populations, is also a genetic locus for sarcoidosis (35). Interestingly, many display the pattern of expression of progressive sarcoidosis being more similar to controls transcripts (also observed at the pathway and upstream regulator level), which may suggest that dysregulation is important in earlier stages of disease onset and then change with progression versus allowing resolution of disease. Because of this, some genes such as CCR5 are upregulated in sarcoidosis compared to controls but downregulated in progressive vs non-progressive comparison. Another plausible explanation for this observation are genetic variants that allow for disease progression, as may be the case for CCR5. These results need further investigation but underscore the importance of analysis by disease phenotypes.

In the non-macrophage population, we identified increased *CCL2* and *TWIST1* in all T cells, which is consistent with previous findings. For example, increased levels of the secreted monocyte chemoattractant CCL2/MCP1 protein have been observed in sarcoidosis BAL (36, 37) and transcription factor TWIST1 (38) in sarcoidosis BAL cells. We also identified activation of T cell exhaustion pathway in CD8+ T cells. In a previous study, exhausted CD8+ T cells were found in sarcoidosis and restoration of CD8+ T cell function led to disease resolution (39).

In the cell-cell interaction analysis, while overall cell interactions were reduced in sarcoidosis, there was a relative increase in CD4+ T cell interactions, indicating a shift in immune dynamics. Key disruptions included downregulation of LGALS9-CD45 signaling, which could impair the stability and function of adaptive regulatory T cell (40). Overall, sarcoidosis is characterized by selective dysregulation in cell communication, particularly involving macrophages and T cells, which may drive immune dysfunction and disease progression.

While the first to evaluate BAL data by phenotypes to this degree, this study has some limitations, including a relatively small sample size, particularly within certain cell subclusters, which may reduce the power to detect significant changes and limit the generalizability of the findings. Interpreting results from pseudobulk analyses can be challenging, as these analyses may mask cell-specific nuances. However, these methods are much more rigorous than earlier studies that likely overestimated changes in gene expression in single cell datasets, including our pilot study (5). As new analytical tools become available, additional analyses will be considered.

In summary, we identified immune cell-specific changes in expression of individual genes, pathways, and upstream regulators associated with disease as well as disease progression. Our cell-cell interaction analysis identified disruptions in communication of antigen-presenting macrophage populations with T cells. Future research should focus on validating these findings and investigating how these genes influence immune dysregulation in sarcoidosis, potentially leading to more precise and effective treatments.

## SUPPLEMENTAL FIGURES

**Supplemental Figure S1.**
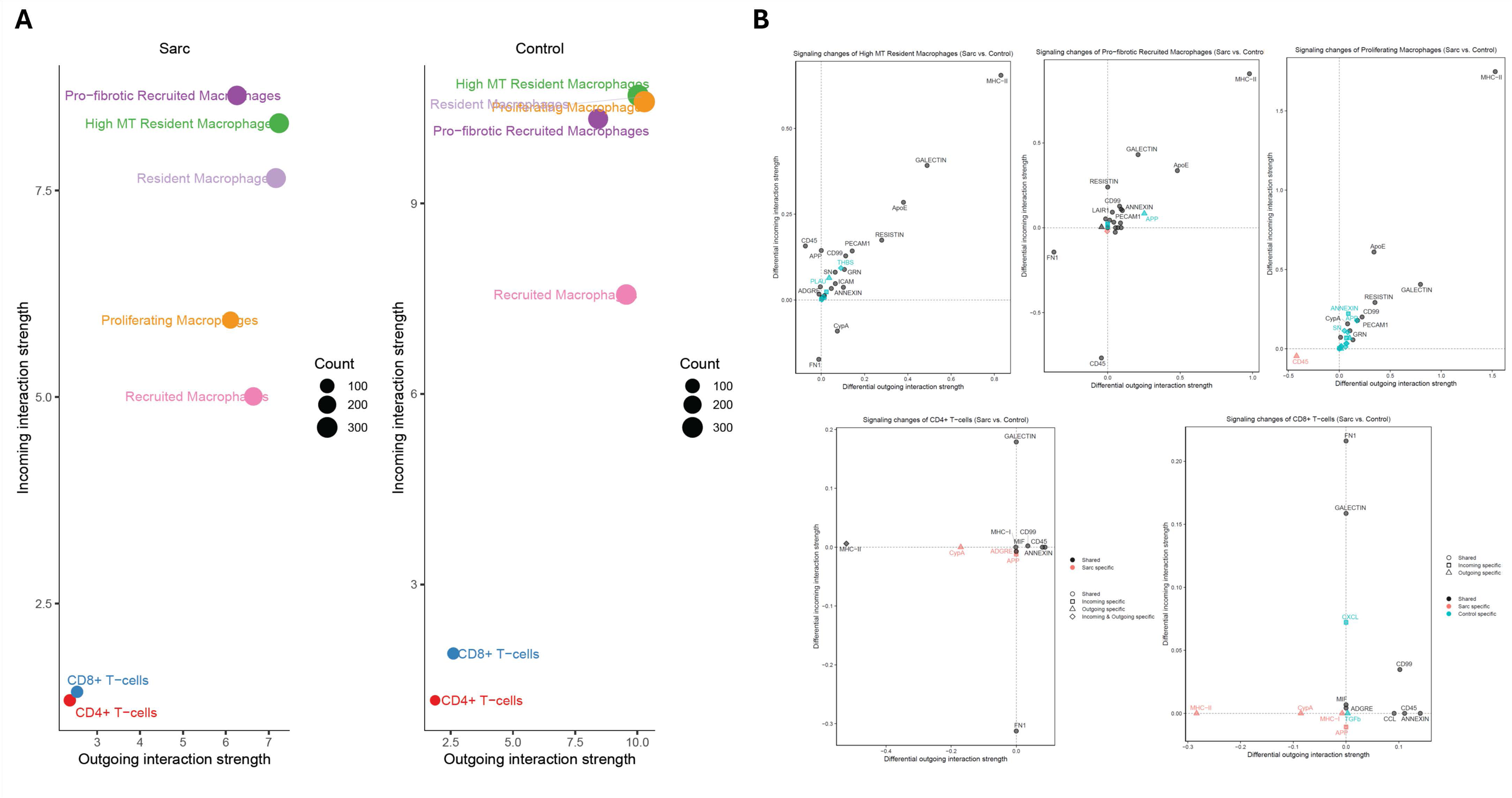
(A) Scatterplots of cell populations with significant changes in sending or receiving signals. **(B)** Scatterplots of differential incoming and outgoing interaction strength for sarcoidosis compared to controls in five cell population (high MT resident macrophages, profibrotic recruited macrophages, proliferating macrophages, CD4+ and CD8+ T cells).

**Supplemental Figure S2.**
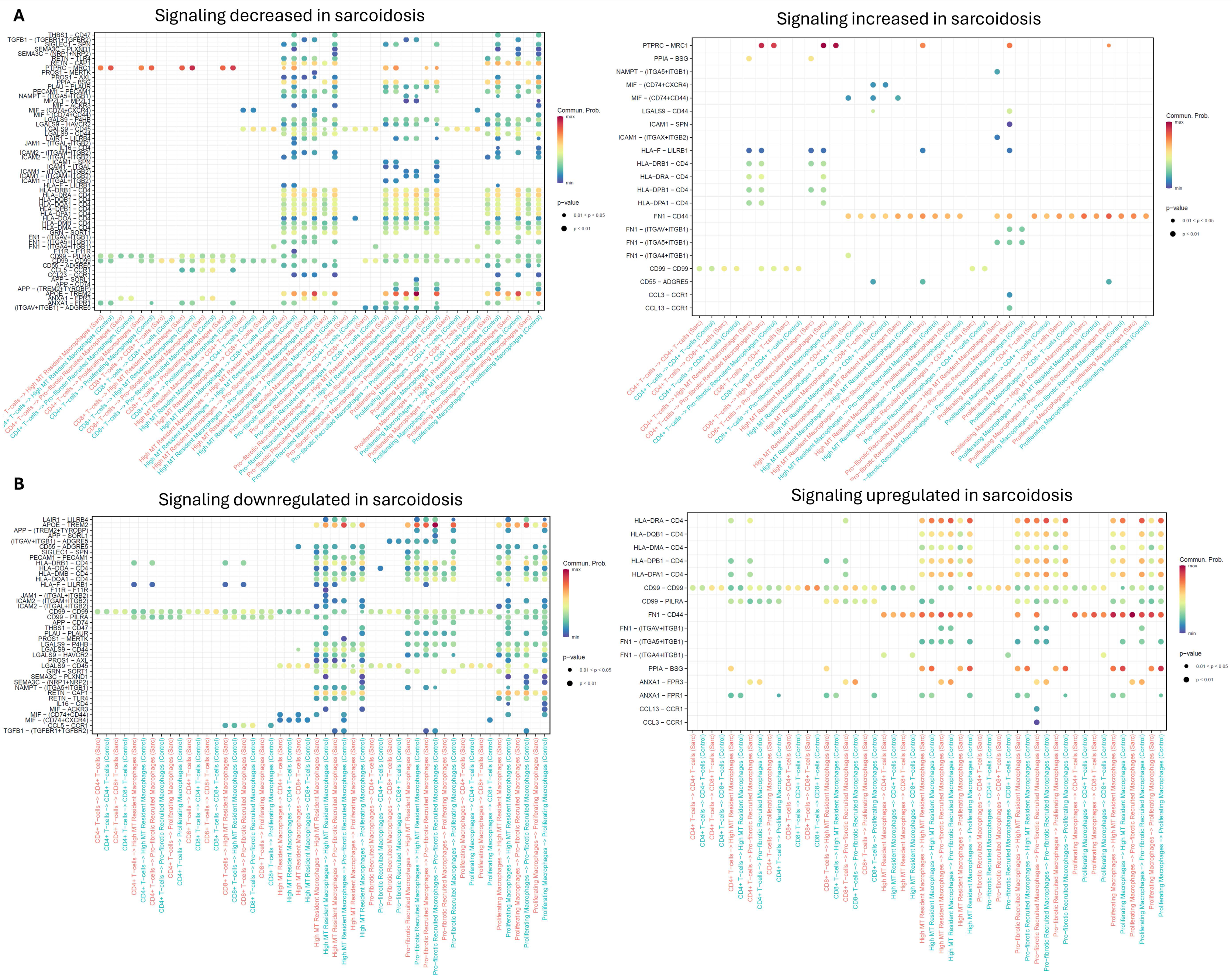
Comparison of specific ligand-receptor interactions in specific cell types that are **(A)** increased or decreased in sarcoidosis, and **(B)** down- and up-regulated in sarcoidosis.

**Supplemental Figure S3.**
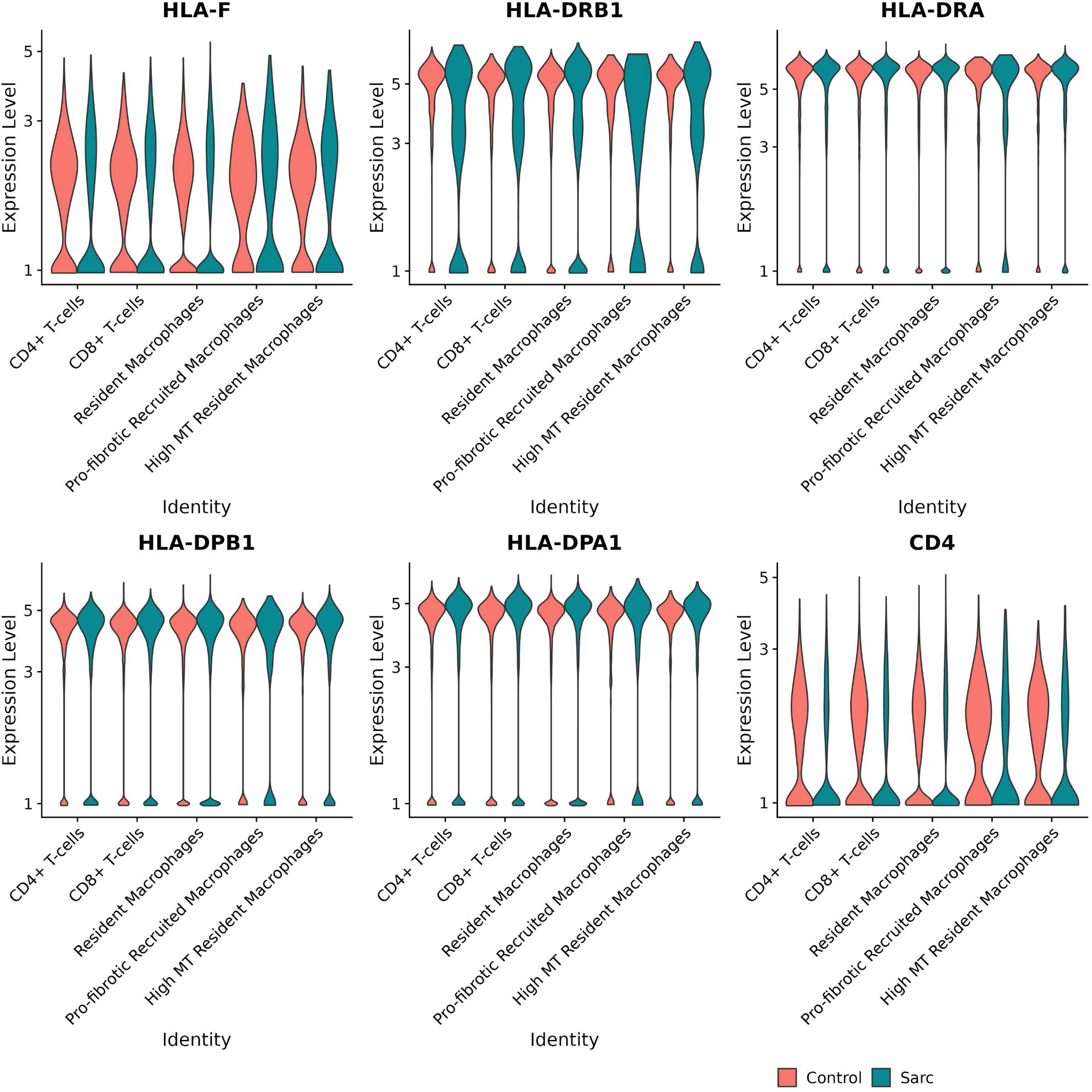
Violin plots of select transcripts identified by CellChat interaction analysis between macrophage and T cell populations.

## SUPPLEMENTAL TABLES

**Supplemental Table S1.** Tests for differences in cell proportions across macrophage clusters in **(A)** sarcoidosis compared to controls and **(B)** progressive sarcoidosis, non-progressive sarcoidosis, and controls

**Supplemental Table S2.** Differentially expressed transcripts in sarcoidosis (N=16) compared to controls (N=14) across macrophage clusters, identified by pseudobulk analysis. **(A)** Resident macrophages, **(B)** high MT resident macrophages, **(C)** recruited macrophages, **(D)** pro-fibrotic recruited macrophages, and **(E)** proliferating macrophages.

**Supplemental Table S3.** Differentially expressed transcripts in sarcoidosis (N=16) compared to controls (N=14) across macrophage clusters, identified by mixed-effects linear modeling of individual cells. **(A)** Resident macrophages, **(B)** high MT resident macrophages, **(C)** recruited macrophages, and **(D)** pro-fibrotic recruited macrophages.

**Supplemental Table S4.** Differentially expressed transcripts in progressive sarcoidosis (N=8) compared to non-progressive disease (N=8) in (A) resident macrophages and (B) recruited macrophages, identified by pseudobulk analysis.

**Supplemental Table S5.** Tests for differences in cell proportions across non-macrophage clusters in **(A)** sarcoidosis compared to controls and **(B)** progressive sarcoidosis, non-progressive sarcoidosis, and controls.

**Supplemental Table S6.** Differentially expressed transcripts in sarcoidosis (N=16) compared to controls (N=14) across non-macrophage clusters, identified by pseudobulk analysis. **(A)** CD4+ T cells, **(B)** CD8+ T cells, **(C)** mixed T cell cluster, **(D)** B cells, **(E)** dendritic cells (DCs), and **(F)** epithelial cells.

## Notes

Supported by NIH awards R01HL140357, R01ES033678, R01ES023826, R01ES025722, R01ES034767 and the Anne Theodore Foundation Breakthrough Sarcoidosis Initiative (ATF-BSI).

### Competing Interest Statement

The authors have declared no competing interest.

